# A Novel Genus of Endogenous Pararetroviruses with Long Terminal Repeats in Grasses

**DOI:** 10.1101/2025.06.08.658526

**Authors:** Dongying Gao

## Abstract

Despite being widespread in plants, endogenous pararetroviruses (EPRVs) are still poorly understood in barley and many other cereal crops. In this study, the barley reference genome was examined and from that a new EPRV was identified and named Hvu-EPRV. In contrast to all EPRVs identified thus far, Hvu-EPRV contains long terminal repeats (LTRs) which are similar to LTR retrotransposons. Homologous sequences of Hvu-EPRV were found in a wide range of plants, however, only those in 17 grasses belonging to the six tribes contain LTRs. The insertion times of nested LTR retrotransposons indicated that Hvu-EPRVs inserted into barley more than 2.37 million years ago, but the invasion and endogenization of Hvu-EPRV related elements in the grass family may be ancient, and horizontal transfers may have occurred between grasses. Phylogenetic analysis revealed that Hvu-EPRV and its homologs in grasses were grouped apart from all 13 reported genera of exogenous and endogenous pararetroviruses, thus the EPRVs in grasses represent a novel genus of the *Caulimoviridae* family named *Moridahovirus*. Genome-wide comparisons of Hvu-EPRVs were conducted between the reference genome and other 84 genomes of cultivated and wild barley, three independent integration events were observed and suggested that the integrations likely occurred after the divergence between barley and its wild progenitor. This is the first time to identify EPRVs with LTRs and to detect their recent integrations, and this research provides new insights into the evolution of plant EPRVs and their invasion history in the grass family.

## Introduction

Plant genomes are complex, with various types of sequences in terms of function and structure. Besides functional genes, plant genomes contain abundant repetitive DNA sequences including both transposons or transposable elements (TEs) and tandem repeats. In some plants, these repeats can contribute over 80% of the genomes (Schnable et al. 2009; Mascher et al. 2017). Once considered ‘junk DNA’, repetitive sequences are now recognized to play essential roles in plant genome evolution, maintenance of centromeric and telomeric stability, and contribute to phenotypic variation (Jiang et al. 2003; Cordaux and Batzer 2009; Gao et al. 2015, Mao et al. 2022). In addition to transposons and tandem repeats, other types of repetitive elements were found in plant genomes including the DNA repeats that share significant sequence identity to the plant pararetroviruses (PRVs).

Pararetroviruses are double-stranded DNA viruses that can infect diverse types of plants and result in serious economic losses. For example, the rice tungro bacilliform virus (RTBV) may induce rice tungro disease and cause annual loss of approximately $1.5 billion or 5-10% of yield reductions in South and Southeast Asia (Dai et al. 2009). However, some plant pararetroviruses also provide useful resources for a broad range of research areas such as the 35S promotor of the cauliflower mosaic virus (CaMV) in plant biotechnology and gene functional analysis. Thus far, 11 genera of exogenous plant pararetroviruses have been identified and all these genera are grouped into the family *Caulimovirida.* The viruses of this family range in size from 7.1 Kb to 9.8 Kb and contain 1–8 open reading frames (ORFs) (Teycheney et al. 2020). Furthermore, plant pararetroviruses exhibit some similar features to mammalian retroviruses as both encode reverse transcriptase (RT) and RNase H (RH) and replicate through an RNA intermediate. However, pararetroviruses do not encode an envelope protein, and integration into the host genome is not obligatory for their life cycle. Instead. they manipulate their episomal copies in the host cell nucleus (Hull and Covey 1996; Geering et al. 2010). As they use reverse transcription for their replication cycle, plant pararetroviruses can be treated as one type of retroelements which also include retroviruses, long terminal repeat (LTR) retrotransposons, non-LTR retrotransposons, and group II mitochondrial introns (Hull 2001; Hansen and Heslop-Harrison 2004). Recently, the family *Caulimoviridae* (pararetroviruses) was grouped together with *Retroviridae* (retroviruses), *Pseudoviridae* (Ty1/Copia LTR retrotransposons), *Metaviridae* (Ty3/Gypsy LTR retrotransposons) and *Belpaoviridae* (Bel/Pao LTR retrotransposons) into the order *Ortervirales* as all these retroelements share a common core Gag-Pol gene (Krupovic et al. 2018).

Endogenous pararetroviruses (EPRVs) are repetitive sequences in plant genomes that show high sequence identity to known exogenous pararetroviruses. They were likely derived from the fortuitous invasions of exogenous pararetroviruses and then have become fixed in plant genomes (Jakowitsch et al. 1999; Staginnus and Richert-Pöggelerde 2006). Except for a few EPRVs in tobacco, petunia, and banana that can be reactivated and become infectious viruses (Lockhart et al. 2000; Richert-Pöggelerde et al. 2003; Tripathi et al. 2019), nearly all the EPRVs identified thus far have degraded and became truncated and dysfunctional sequences, thus these complete EPRVs were reconstructed with multiple homologous sequences (Jakowitsch et al. 1999; Geering et al. 2014). Since first discovery in tobacco (Jakowitsch et al. 1999), EPRVs were detected in various plants including both angiosperms and gymnosperms (de Tomás and Vicient 2022; Vassilieff et al. 2023). Phylogenetic analysis indicated that some EPRVs were separated from the reported PRVs and suggested that they may represent new genera of the *Caulimoviridae* family (Geering et al. 2014; de Tomás and Vicient 2022). The integration of EPRVs may potentially affect gene expression and cause deleterious mutations. However, they can also bring benefits to the hosts such as the control of related exogenous pararetroviruses via RNA silencing pathways (Hohn 2015; Valli et al. 2023). Furthermore, EPRVs are genomic fossils and provide insights into the evolutionary history of PRVs’ infections and endogenizations in the host genomes (Gayral et al. 2010).

Barley (*Hordeum vulgare*, 2n=2x=14) is an important crop widely consumed for food, animal feed and malting industries in the world. It is a diploid species and shares close evolutionary relationships with several polyploid cereals such as wheat (*Triticum aestivum*) and oat (*Avena sativa*), thus barley can be used as a model organism for cereal genomics. Thus far, nearly 90 genomes of cultivated barley and its wild progenitor (*Hordeum spontaneum*) have been sequenced and analyzed (Mascher et al. 2017; Liu et al. 2020; Zeng et al. 2020; Sato et al. 2021; Xu et al. 2021; Jiang et al. 2022; Hu et al. 2023; Pan et al. 2023; Clare et al. 2024; Jayakodi et al. 2024). However, only fragmentary EPRVs were identified in barley (de Tomás and Vicient 2022). In this study, the barley reference genome was investigated and a new EPRV which contains LTR was identified and named Hvu-EPRV. The phylogenetic analysis suggested that the EPRV may belong to a novel genus of the family *Caulimovirida*. The barley pan-genomic sequences were compared to reveal polymorphic EPRVs among the sequenced barley genomes suggesting the recent integrations of Hvu-EPRVs.

## Results

### A New Endogenous Pararetrovirus with Long Terminal Repeats in Barley

To detect endogenous pararetrovirus sequences in barley, the conserved domains of reverse transcriptase (RT) proteins of 22 reported plant pararetroviruses (Vassilieff et al. 2022) were used to search against the barley reference genome-Morex V3 (Mascher et al. 2017). Numerous significant hits were detected, and the lowest E-value was 3.4 × e^-107^. As other retroelements including both LTR retrotransposons and long interspersed nuclear elements (LINEs) also contain the RT domain, to confirm the presence of EPRVs and determine their boundaries, the 20-Kb flanking sequences of the hits with very lower E-values (< 1 x e^-100^) were extracted and used to search against the barley genome. After manual curations and pairwise sequence comparisons, a novel endogenous pararetrovirus called Hvu-EPRV was identified. The size of the representative element was 9,446 bp and it was located on Chromosome 2H (GenBank Accession: LR890097: 8906251-8896806). This element contains two ORFs, the first one encodes a 1,748-aa protein which contains the domains of reverse transcription, RNase H (RH), GAG/COAT and movement protein (MP) (**Fig. 1a**), and the second one, ORF2, encodes a 168-aa protein. The function of the ORF2 is not clear as it did not share significant sequence homology to any known proteins or domains, and the only weakly related hit was the transactivator/viroplasmin (TAV) domain of the soybean chlorotic mottle virus (X15828, E-value = 0.42).

**Figure 1.**
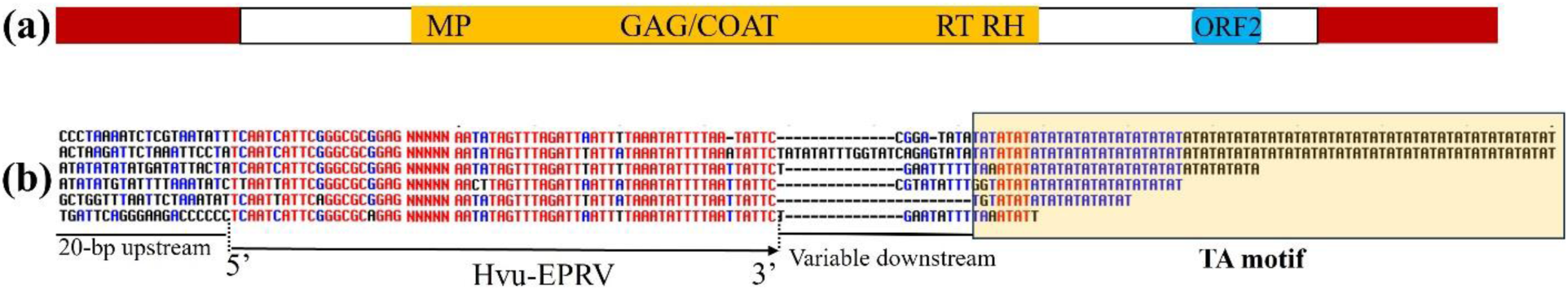
a. Sequence structure of Hvu-EPRV element. The brown rectangles indicate the long terminal repeats, **b. Alignment of six Hvu-EPRVs and their flanking sequences.** The internal regions are represented by “NNNNN”, and the TA simple sequence repeats are boxed.

Close evolutionary relationships were revealed between plant pararetroviruses and LTR retrotransposons (Boeke 2003; Hansen and Heslop-Harrison 2004; Krupovic et al. 2018). However, none of the reported plant pararetroviruses including both exogenous viruses and the endogenous elements has LTR (Richert-Pöggeler et al. 2003; Staginnus and Richert-Pöggelerde 2006; Tomás and Vicient 2022; Vassilieff et al. 2022). Like many EPRVs, Hvu-EPRV also lacks the integrase (IN) domain, but it contains LTRs which are similar to the termini of LTR retrotransposons. The LTRs of Hvu-EPRV was about 1,300 bp, they start with ‘TCAAT’ and end with ‘TATTC’ motif (**Fig. 1b**) which are different from the terminal motifs of LTR retrotransposons (usually 5’ TGT…ACA3’). Also, no target side duplications (TSDs) were found for all Hvu-EPRV copies, the common feature for the flanking sequences at their 3’ end was the variable units of “TA” motif. Therefore, Hvu-EPRV represents an unusual endogenous pararetrovirus. This is the first time a plant pararetrovirus with LTRs has been identified.

### Tandem Organizations and Nested Insertions

To detect its abundance and distributions, the representative element of Hvu-EPRV was used to screen the barley reference genome. A total of 383 hits were detected which have masked 956,316-bp or 0.02 % of the barley reference genome (4,195,169,932 bp, gaps were ignored). As some hits were short (about 50-bp) and a single Hvu-EPRV element can match multiple hits. Therefore, manual inspections were conducted, 271 Hvu-EPRV elements with high confidence (the Smith-Waterman score > 1000 and the hit sizes > 150 bp) were retained for further analysis. These elements were distributed on all seven barley chromosomes, the Chromosome 7H contains the most abundant elements (71 Hvu-EPRVs) whereas the Chromosome 5H harbors the lowest number of sequences (19 Hvu-EPRVs). Most of the Hvu-EPRV elements were dispersed in the barley genome, but tandem organizations were also found for which multiple Hvu-EPRV elements are repeated in a head-to-tail manner (**Fig. 2a-c**). Additionally, some Hvu-EPRV elements are nested by one or more LTR retrotransposons (**Fig. 2d-e**) or other transposons. As LTR retrotransposons replicate through reverse transcription and their LTR sequences were identical when they inserted, thus the insertion times of LTR retrotransposons can be estimated based on the sequence divergence of the two LTRs (Perlman and Boeke 2004). Six intact LTR retrotransposons nested by Hvu-EPRVs were identified, and their insertion times ranged from 169,231 to 2,365,385 years ago (Table 1). It also suggested that Hvu-EPRV elements have already been integrated into the barley genome more than 2.37 million years ago (Mya) because the insertions of nested LTR retrotransposons should be later than the target Hvu-EPRV sequences.

**Figure 2.**
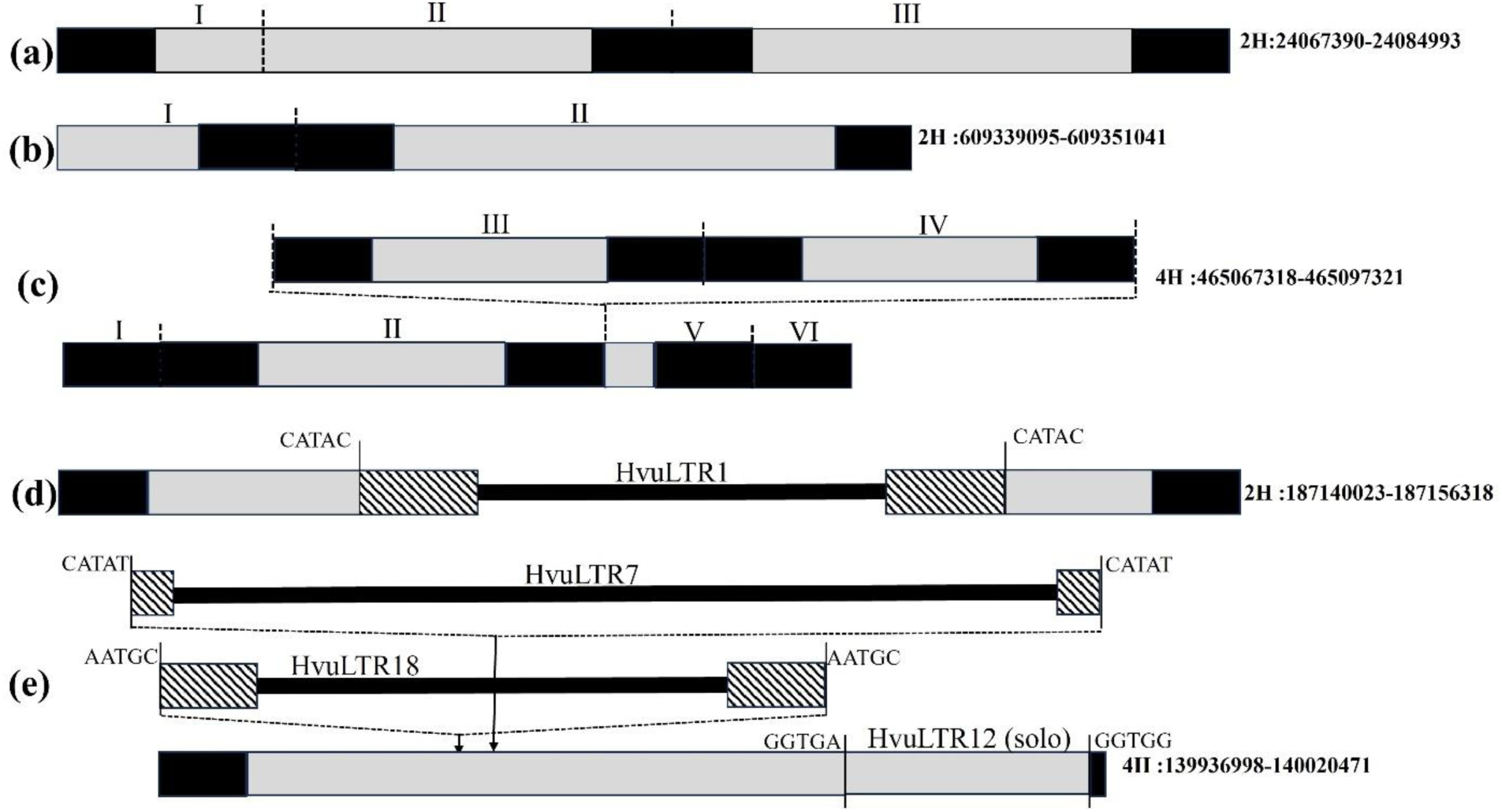
a-c. Tandem organizations of Hvu-EPRV sequences. d-e. Hvu-EPRVs nested by LTR retrotransposons. The black and grey rectangles mean LTRs and internal regions of Hvu-EPRVs. The different copies of Hvu-EPRVs are indicated by Roman numerals which were defined based on the 9,446-bp reference element. The striped rectangles and thick black lines represent the long terminal repeats and the internal regions of retrotransposons. The vertical black lines indicate the target site duplications.

**Table 1.** Estimation of six full-length LTR retrotransposons nested by barley pararetroviruses

| Chromosome | Position of Hvu-EPRV | Position of nested LTR | Name/Superfamily of LTR retrotransposon | Target site duplication | Insertion time (Year) |
| --- | --- | --- | --- | --- | --- |
| LR890097 (2H) | 187140023-187156398 | 187144110-187153050 | Hvu-LTR1/Copia | CATAC | 169,231 |
| LR890099 (4H) | 139936998-140020471 | 139941031-139950242 | Hvu-LTR7/Gypsy | AATGC | 1,942,308 |
|  | 139936998-140020471 | 139950498-139963905 | Hvu-LTR18/Gypsy | CATAT | 557,692 |
|  | 139936998-140020471 | 139973098-139981708 | Hvu-LTR1/Copia | CTAGG | 255,385 |
|  | 139936998-140020471 | 140007001-140015254 | Hvu-LTR1/Copia | CAAGG | 2,365,385 |
| LR890102 (7H) | 66389911-66409985 | 66395136-66409559 | Hvu-LTR39/Gypsy | TATAT | 1,007,692 |

Note: The Morex Version 3 (V3) was used for this analysis.

Tandem EPRVs have been reported in other plants (Richert-Pöggeler et al. 2003; de Tomás and Vicient 2022), given that the barley genome is highly repetitive and contains numerous highly identical and large transposons which may affect correct genome assemblies (Phillippy et al. 2008). To further confirm the tandem arrangements of Hvu-EPRV, the 9,446-bp Hvu-EPRV element was used to screen the long-read sequencing data (10.5 Gb or 531,066 reads) which was previously generated by a Nanopore GridION sequencer for genomic analysis of a transgenic barley cultivar (Gao et al. 2024). After manual curations, 409 unique reads were found to contain Hvu-EPRV sequences. Among these long reads, 111 were tandemly repeated and contain two or more Hvu-EPRV copies (**Supplementary Fig. S1a-d**). Therefore, the long-read sequencing data further confirmed tandem organizations of Hvu-EPRV in barley genome. The long-read sequencing data also confirmed that Hvu-EPRVs contain LTRs.

### Homologous Sequences of Hvu-EPRVs are Present in A Wide Range of Plants

Hvu-EPRV was used to search against the sequenced plant genomes deposited in GenBank and identify its homologous elements. Homologous sequences of Hvu-EPRV were identified in 81 plant species including five gymnosperms (*Cunninghamia lanceolata*, *Taxus chinensis*, *Pinus lambertiana*, *Ginkgo biloba* and *Gnetum montanum*), two ferns (*Polystichum acrostichoides* and *Dryopteris marginalis*) and 74 flowering plants which contained 45 monocots, 26 dicots, two magnoliids, and the ancient angiosperm *Nymphaea thermarum* (**Supplementary Table S1**). It should be noted that the homologs of Hvu-EPRVs may also be present in more plant genera in GenBank as very strict criteria were employed to define the homologous sequences (E-value < 1 × e^-10^ and the hit sizes > 1,000 bp). Impressively, the homologous sequences in 17 grass plants contain LTRs and all of these plants belong to the six tribes of the grass family (Poaceae) including Triticeae, Poeae, Brachypodieae, Bambuseae, Eragrostideae, and Cynodonteae (**Supplementary Table S1**). Pairwise sequence comparisons with Hvu-EPRVs indicated that both terminal repeats and the internal regions (protein-coding) of these elements showed higher sequence similarity to that of Hvu-EPRV, but the internal regions seemed more conserved (**Supplementary Fig. 2**). The homologs in other grass tribes and other plant families also shared significant sequence identity, but they lack repetitive termini. Additionally, no significant hit was found in some plants including *Arabidopsis thaliana*, maize (*Zea mays*) and sorghum (*Sorghum bicolor*) suggesting that these plants may never be invaded by pararetroviruses or the EPRVs in these plant genomes have been eliminated by the genomic pressures.

### Phylogenetic analysis of Hvu-EPRV

To understand the evolutionary relationships between Hvu-EPRV and other plant EPRVs, the RT conserved domains of 43 plant EPRVs identified in this study which names ended with “-EPRV” (**Supplementary Table S1**) was used to conduct phylogenetic analysis. Thirty-nine other identified plant EPRVs including the EPRV in wheat (Tae-EPRV) were excluded as they lack complete RT conserved domains. Furthermore, the RT conserved domains of 37 previously described retroelements were also included for investigating the phylogenetic evolution of plant EPRVs, which included four Ty3/Gypsy LTR retrotransposons (*Metaviridae*) and 33 exogenous and endogenous pararetroviruses which represent all 13 genera of the family *Caulimovirida* reported thus far (**Supplementary Table S2**). The four LTR retrotransposons used as the outgroup were grouped into clade I, and the 76 plant pararetroviruses were grouped into Clade II to Clade XVI (**Fig. 3**). Florendovirus_Nsyl and CitIch-033, belonging to the two new genera of *Caulimovirida*, florendovirus (Geering et al. 2014) and Wendovirus (de Tomás and Vicient 2022), was grouped into clade III and VI, respectively. Impressively, Hvu-EPRV was grouped apart from all the 33 reported pararetroviruses but it was grouped together with the homologous EPRVs identified in other grass genomes, and all these 15 EPRVs including the EPRVs with LTRs fell into the Clade VII. Thus, the phylogenetic analysis suggested that Hvu-EPRV and other EPRVs in grasses likely represent a novel lineage of the family *Caulimovirida*.

**Figure 3.**
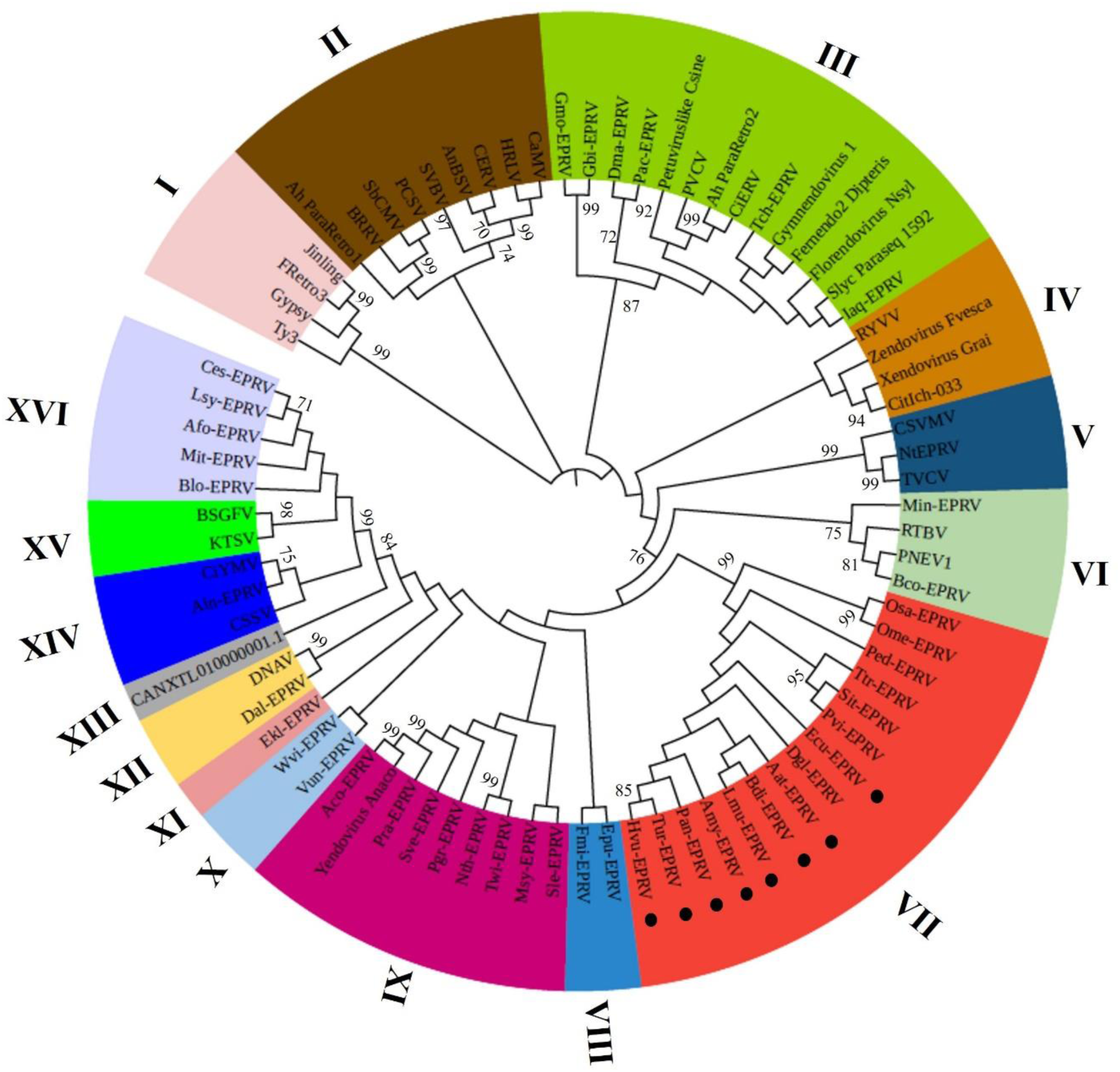
Phylogenetic tree of RT conserved domains of four Ty3/Gypsy LTR retrotransposons and 76 exogenous and endogenous pararetroviruses. The 43 EPRVs identified in this study end with “-EPRVs” and the black ovals indicate that the EPRVs contain LTRs. The bootstrap values of > 70% are labeled. See Table S1 and Table S2 for detailed information about sequences used in the analysis.

### Evolution of EPRVs in the Grass Family

Except the canonical telomeric tandem repeats and few transposon families, most repeats in the plant kingdom evolved rapidly and homologous repetitive repeats were difficult to observe between related species (Jiang et al. 2003; Gao et al. 2009). It was unexpected that significant sequence similarity between the EPRVs in different grasses was detected at the nucleotide level, such as 88% sequence identity between Hvu-EPRV and Tae-EPRV in wheat and 78% identity between Ped-EPRV in *Phyllostachys edulis* and Pau-EPRV in *Phragmites australis*. To test the possibility of horizontal transfers of EPRVs, the EPRVs in 33 grasses belonging to 17 tribes were used to conduct phylogenetic analysis. As LTRs were more divergent than the internal coding regions (**Supplementary Fig. S2**) and not all EPRVs in grasses contain LTRs, thus only the internal coding DNA sequences of the EPRVS were used to construct the phylogenetic tree. The EPRVs from the three non-grass plants (outgroup), including *Dioscorea alata*, *Aristolochia californica* and *Annona cherimola*, were grouped into the clade X, and all EPRVs in grasses fell into nine clades (I to IX) (**Fig. 4**). In most cases, the EPRVs from same plant genomes and tribes were grouped together which is consistent with the evolutionary origins of the grasses. However, phylogenetic incongruence was also found. For example, the EPRVs in *Brachypodium distachyon* were grouped together with the EPRVs in the Poeae tribe but not separated from the Poeae and Triticeae tribes. Also, the EPRVs in common reed (*Phragmites australis*, the Molinieae tribe of PACMAD clade) were grouped into clade VII, together with the EPRVs in the three tribes of BOP clade including *Dendrocalamus latiflorus* (Bambuseae), *Stipa capillata* (Stipeae), and *Phyllostachys edulis* (Arundinarieae). Thus, the phylogenetic analysis implied the potential horizontal transfers of EPRVs in grasses.

**Figure 4.**
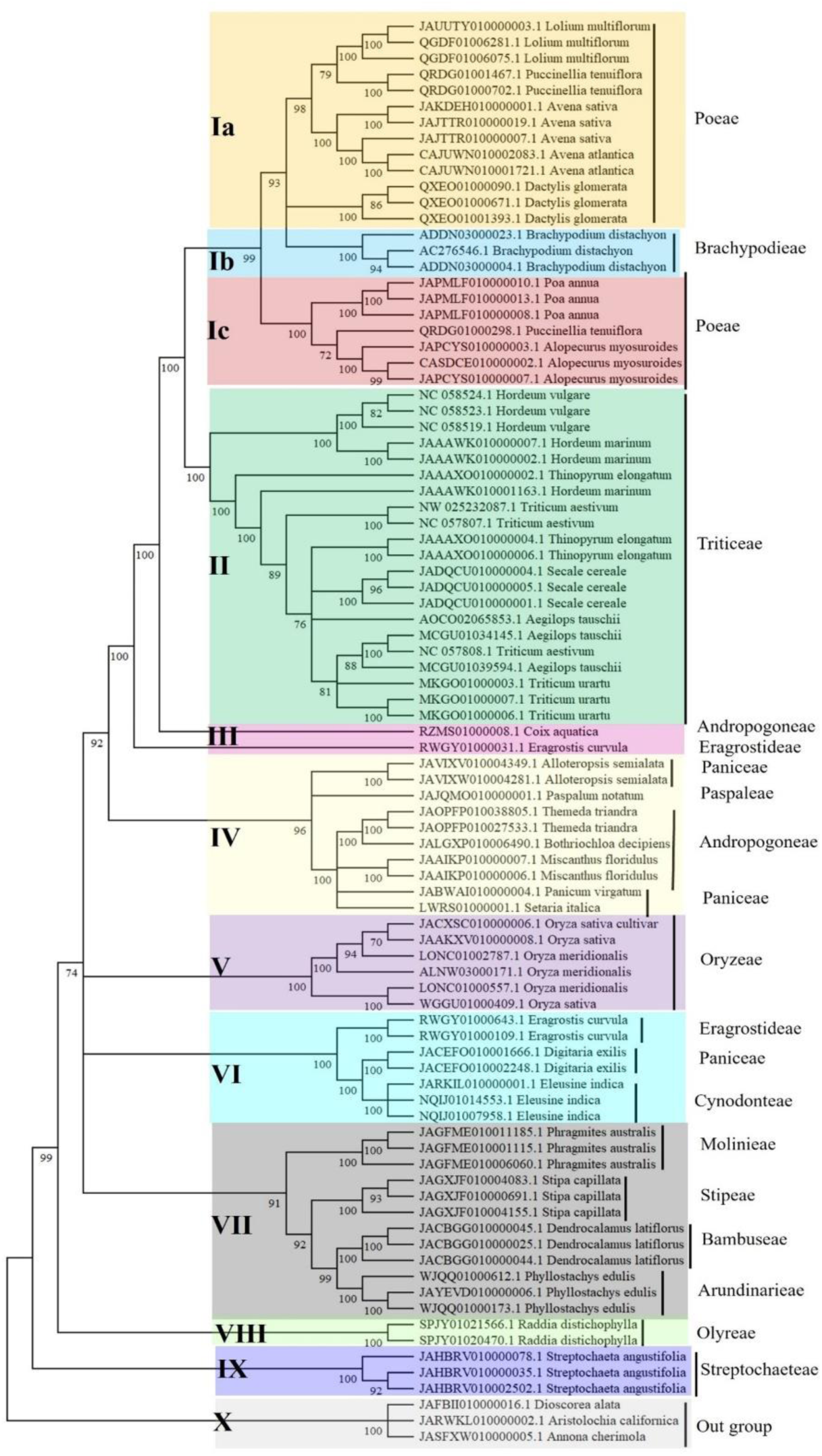
The phylogenetic trees constructed with the internal DNA sequences of EPRVs in 33 grasses and three non-grasses (outgroup) including *Dioscorea alata*, *Aristolochia californica* and *Annona cherimola*. The bootstrap values of > 70% are labeled.

### Barley Pan-Genomes Revealed Recent Integrations of Hvu-EPRVs

During sequence comparisons, multiple hits in the barley reference genome demonstrated high sequence identity (99% for over 9.4-Kb aligned sequence size) to the 9,446-bp representative Hvu-EPRV element. To provide more evidence on the recent activity of Hvu-EPRVs, genome-wide comparisons of Hvu-EPRVs were further conducted. As of December 26, 2024, a total of 197 genomes of cultivated and wild barley have been deposited in GenBank. After removing the redundant genomes (same genotypes but different versions of submissions), 85 published genomes with high quality were used for the comparative analysis which include 61 cultivated barley and 24 wild species (*H. spontaneum*) (**Supplementary Table S3**). Among the 271 Hvu-EPRVs with high confidence identified in Morex, most of them were significantly truncated, but 29 Hvu-EPRVs are intact or nearly complete (200-bp shorter than the representative Hvu-EPRV sequence) that may imply their recent integrations. The 2-Kb (1-Kb for each side) merged flanking sequences of the Hvu-EPRVs were extracted from the Morex genome and used to search against the 84 genomes of cultivated and wild barley. Polymorphic Hvu-EPRVs were identified at eight genomic loci (**Fig. 5a**, **Supplementary Table S3**) including three positions (II, III and IV) where Hvu-EPRVs are absent in all 24 wild barley genomes, thus they likely inserted into the barley genomes after the split of cultivated and wild barley. The Hvu-EPRVs at other five loci (I, V, VI, VII, and VIII) are shared between cultivated barley and the wild species (**Supplementary Table S3**) implying the integrations of these Hvu-EPRVs likely occurred before the divergence. To gain insights into the integrations of Hvu-EPRVs, the immediate flanking sequences of the polymorphic Hvu-EPRVs in Morex and the other ten cultivated and wild barley genomes were aligned, TA simple sequence repeats were found for both the upstream and downstream flanking regions (**Fig. 5b, Supplementary Fig. 3a, c**). In addition, TG tandem repeat was also detected for the downstream flanking region of the Hvu-EPRVs (**Supplementary Fig. 3b**). These results suggested that Hvu-EPRVs prefer to integrate into the TA or TA-TG tandem repeats.

**Figure 5.**
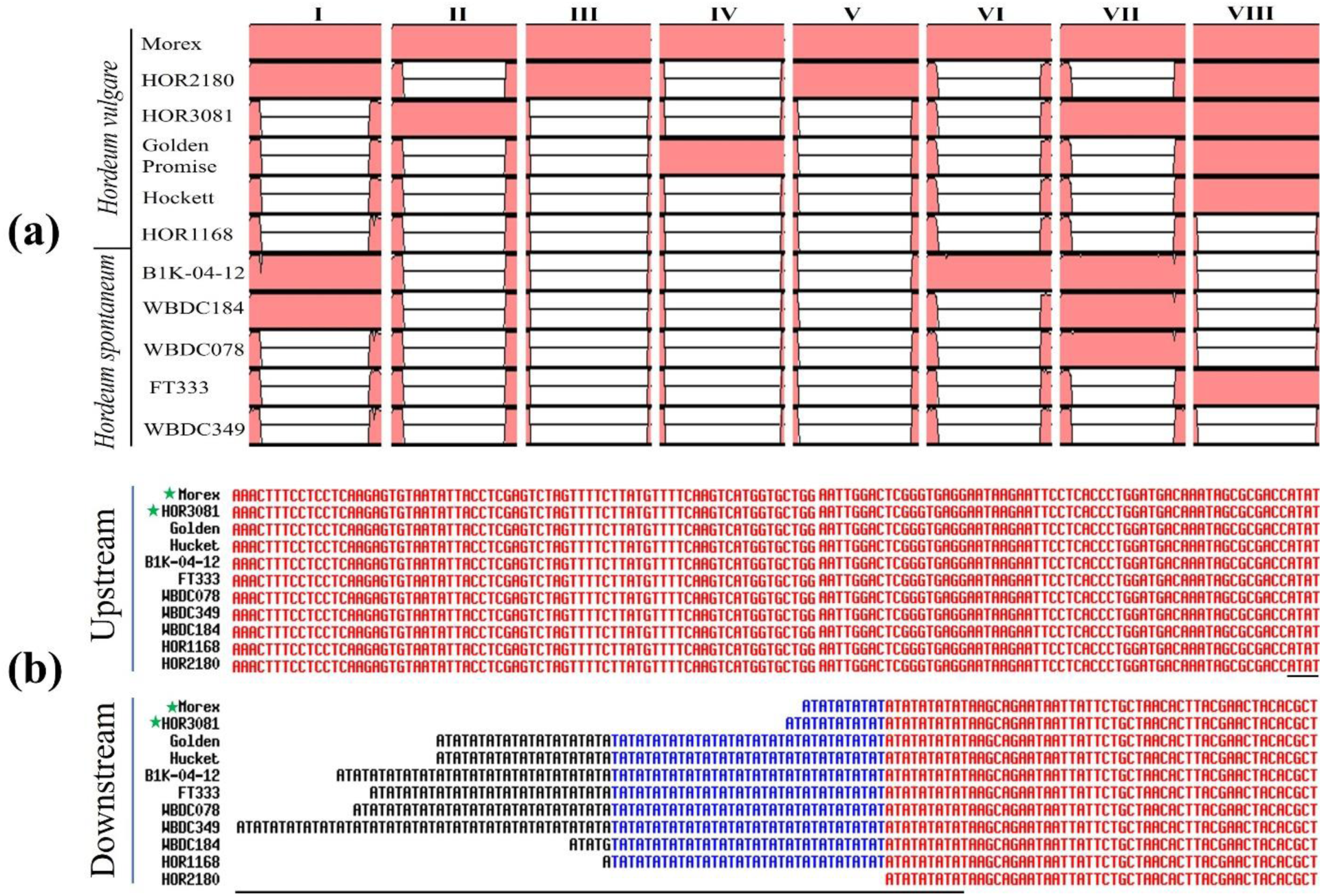
a. Comparisons of Hvu-EPRVs and the flanking sequences (1-Kb for each side) between Morex and 10 cultivated and wild barley genomes. b. Alignment of 5’ (up) and 3’ (down) sequences immediately flanked the Hvu-EPRV in Morex (LR890098: 345974868-345984153) and 10 cultivated and wild barley genomes. The TA simple repeats are underlined. The green stars mean the genotypes containing Hvu-EPRV.

## Discussion

### A New Genus of the *Caulimovirida* Family

Thus far, nearly 90 genomes of cultivated and wild barley have been sequenced and analyzed (Mascher et al. 2017; Liu et al. 2020; Zeng et al. 2020; Sato et al. 2021; Xu et al. 2021; Jiang et al. 2022; Hu et al. 2023; Pan et al. 2023; Clare et al. 2024; Jayakodi et al. 2024). However, only fragmentary EPRVs (less than 1,000 bp) were reported in barley (de Tomás and Vicient 2022). In addition, no EPRV was identified in many other published grass genomes including the major cereal crops such as wheat (International Wheat Genome Sequencing Consortium, 2018), rye (*Secale cereale*) (Li et al. 2021) and oat (*Avena sativa*) (Kamal et al. 2022). In this study, the barley reference genome was analyzed and a new EPRV named Hvu-EPRV was identified. Further sequence searches against the barley pan genomes and other plant genomes indicated that EPRVs are also present in wild barley, wheat, rye, oat, *Thinopyrum elongatum* and other grasses (**Supplementary Table S1**). EPRVs have been found in a wide range of plants including both angiosperms and gymnosperms (Jakowitsch et al. 1999; de Tomás and Vicient 2022; Vassilieff et al. 2023), and all identified EPRVs and exogenous pararetrovisuses lack LTRs (Richert-Pöggeler et al. 2003; Staginnus and Richert-Pöggelerde 2006; Geering et al. 2014; Tomás and Vicient 2022; Vassilieff et al. 2022). However, Hvu-EPRV and its homologous elements in 17 grasses do contain LTRs (**Supplementary Table S1**), although these LTRs do not have the classical “5’TGT…ACA3’” terminal motifs of most typical LTR retrotransposons. This is the first time EPRVs with LTRs have been identified. The phylogenetic analysis of RT conserved domains indicated that Hvu-EPRV and its homologous sequences in grasses were grouped apart from all 13 known genera of the *Caulimovirida* family (**Fig. 3**). Based on their unique sequence structures and the phylogenetic split, Hvu-EPRV and other EPRVs in grass genomes may represent a new genus of the *Caulimovirida* family named *Moridahovirus*. It should be noted that the homologous sequences of Hvu-EPRV in other plants may belong to the identified genera or other new genera that may be defined in the future. For example, the homologs of Hvu-EPRV in *Ananas comosus* (Aco-EPRV) and *Puya raimondii* (Pra-EPRV) were grouped together with Yendovirus_Anaco belonging to the *Yendovirus* genus (Vassilieff et al. 2022) into the clade IX and all the EPRVs in this clade lack LTRs.

### Evolutionary origin of the EPRV with LTRs

Plant pararetroviruses are retroelements, they are considered as the sister lineage of Ty3/Gypsy LTR retrotransposons or were likely originated from Ty3/Gypsy LTR retrotransposons (Boeke 2003; Hansen and Heslop-Harrison 2004; Krupovic et al. 2018). Therefore, pararetroviruses have undergone at least two important events during evolutionary time: 1) loss of LTRs and integrase domain, and 2) acquisition of the related proteins for virus transmission and infection. The homologous sequences of Hvu-EPRV were found in 77 plants spanning flowering plants, gymnosperms and ferns (**Supplementary Table S1**), this widespread presence suggested the ancient origin of EPRVs in plants. However, the homologs of Hvu-EPRV in non-grasses were either grouped together with the previous genera of pararetroviruses or grouped into different clades (**Fig. 3**). Additionally, all EPRVs identified in the non-grasses lack terminal repeats. Therefore, the EPRVs with LTRs likely emerged after the split between the grass (Poaceae) and other plant families. In this study, EPRVs with LTRs were identified in 17 grasses belonging to the four tribes of the BOP clade (Triticeae, Poeae, Brachypodieae and Bambuseae) and two tribes of PACMAD clade (Cynodonteae and Eragrostideae). The BOP and PACMAD clades were diverged from each other about ∼73.6 million years ago (Mya) (Ma et al. 2021), it was possible that the acquisition of LTRs occurred before the divergence of BOP clade and PACMAD clade. It should be noted that the EPRVs in many grasses, including rice (*Oriza sative*, the Oryzeae tribe of the BOP clade) and switchgrass (*Panicum virgatum*, the Paniceae tribe of the PACMAD clade), also lack LTRs. One explanation for this may be that the LTRs in these EPRVs have been lost or removed. Another scenario was that EPRVs in one or few grasses acquired LTRs after the divergence between the BOP and PACMAD clades and then horizontal transfers occurred between the grasses as the phylogenetic incongruence of EPRVs was also found between some grasses (**Fig**. **4**) LTRs do not encode proteins, but they contain regulatory elements including enhancers, promoters, and termination signals that are important for the replication and integration of LTR retrotransposons (Temin 1981; Chang and Schulman 2008). As LTR retrotransposons represent the most abundant repetitive sequences in many, may be all, sequenced plant genomes (Schnable et al. 2009; Mascher et al. 2017; Jayakodi et al. 2024), it was possible that LTRs may also provide some beneficial impacts on amplification and burst of LTR retrotransposons in plants. Sequence search against the database of plant cis-acting regulatory DNA elements (Higo et al. 1999) identified several regulatory motifs including TATA box (TATTTAA) in the LTR region of Hvu-EPRV, it is not clear whether the LTRs are involved in and benefit the replication of Hvu-EPRV and its homologs in grasses. Despite the evolutionary origin and function of the LTRs of EPRVs are still needed to be investigated with more analysis and experiments, Hvu-EPRV and its homologous elements in grasses provide new insights into the evolution of plant pararetroviruses as they belong to a unique lineage of pararetroviruses and may represent the intermediate or transition stage from pararetroviruses that lack LTRs to LTR retrotransposons.

### Recent activity of Hvu-EPRV

Recently, de Tomás and Vicient (2022) analyzed 278 plant genomes, conducted sequence comparisons between different copies of a same EPRV and detected 100% sequence identity at the RT conserved regions of some EPRVs. In this study, high sequence identity (99% over 9-Kb aligned regions) was detected between different copies of Hvu-EPRV, and further genome-wide comparisons between Morex and other 84 sequenced cultivated and wild barley identified eight polymorphic Hvu-EPRVs. Impressively, three of them are only present in some cultivated barley but absent in all wild barley genomes (**Supplementary Table S3**). Thus, the comparisons of barley pangenomes indicated that the integrations of some Hvu-EPRVs were recent, and the endogenization events even occurred after the divergence between cultivated barley and its wild progenitor. In this study, only three independent recent integrations of Hvu-EPRVs were identified, it is expected that there are more recent insertions of Hvu-EPRVs in the natural barley population including some that may be specific to wild barley or the cultivated barley of the new continents because only the complete elements in Morex were compared with other barley genomes.

Except the EPRVs in tobacco, petunia and banana which may still contain replication competent sequences and can be reactivated to infect plants under certain conditions (Lockhart et al. 2000; Richert-Pöggelerde et al. 2003; Tripathi et al. 2019), the endogenizations of most plant EPRVs were very ancient as they are heavily degraded and transcriptionally inactive (Jakowitsch et al. 1999; Geering et al. 2014). However, Hvu-EPRVs seem exceptional as high sequence similarities were detected between multiple full-length copies. In addition, 88 expressed sequence tags (ESTs) from barley, wheat and other 15 grasses showed significant sequence identity to Hvu-EPRV (E-value < 1 x e^-10^) were detected (**Supplementary Table S4**) that suggested the possible expression of Hvu-EPRV and its homologs.

Like other retroelements, EPRVs can insert into genes and disrupt their functions. However, the integrations of endogenous viruses may be beneficial and will be fixed by the hosts (Isbel and Whitelaw 2012). The EPRVs in grasses may also contribute to gene content. For example, Tae-EPRV was used to search the wheat transcriptional data sets (Ramírez-González et al. 2018) and detected one gene, *TraesCS2A02G092000*, that exhibited significant (4.2 x e^-58^) sequence identity. The gene encodes cysteine-rich receptor kinase (CRK) protein and contains partial LTR sequence of Tae-EPRV (**Supplementary Fig. S4a**). The gene *TraesCS2A02G092000* was expressive under stress treatments including the disease infections (**Supplementary Fig. S4b**). Previous research indicated that EPRVs are frequently inserted into the TA-rich regions which are associated with chromosome fragility (Geering et al. 2014). In this study, TA simple sequence repeats were frequently found in the flanking regions of Hvu-EPRVs, but TG simple sequence repeats also surround some Hvu-EPRVs (**Fig. 4b, Supplementary Fig. 3**) which may indicate the insertion preference of Hvu-EPRVs or the sequence capture of double-stranded DNA break repair. Despite transcriptional activity of Hvu-EPRVs was detected by the sequence searches, it seems that Hvu-EPRV has lost its infection ability as its TAV domain (ORF2) was truncated and highly divergent. Furthermore, no significant hit was found in the barley genome when the TAV protein was used to conduct TBLASTN search. It was reported that TAV domain plays a critical role for the basic replication of pararetrovirus (Kobayashi and Hohn 2003). The TAV mutations of Hvu-EPRV should impact its infection and transmission although more experiments are needed to be conducted in the future.

In conclusion, this study identifies new EPRVs with long terminal repeats in barley and other grass genomes. Given their unique structures and phylogenetic segregation, the EPRVs in grasses represent a new genus of pararetrovirus called *Moridahovirus*. The EPRVs may have inserted into the barley genome more than 2.37 million years ago, but their origin in grass may be ancient and the horizontal transfer of EPRVs cannot be excluded. Some EPRVs in barley are complete and may integrated into the barley genome after the divergence with wild species. Overall, this is the first time to identify EPRVs with LTRs and reveal their recent integration, the results provide new insights into the evolution and function of plant pararetrovisuses.

## Methods

### Identification of Endogenous Pararetroviruses in Barley and Other Plants

To identify endogenous pararetrovirus sequences in barley, the RT proteins of different plant pararetrovirus genera (Vassilieff et al. 2022) were used to conduct TBLASTN searches against the barley reference genome (Mascher et al. 2017). The flaking sequences (10-Kb for each side) of the significant hits (E-value < 1 × e^−6^) were extracted and merged with the hits and used to search against the barley genomic sequences for defining the boundaries of the endogenous pararetroviruses in barley. The representative barley pararetrovirus was then used to search against GenBank and find the homologous elements in other plants and their boundaries were further determined by comparing the homologous sequences and their flanking regions.

To predict the structures of plant pararetroviruses and define their domains, the pararetroviruses were first annotated by GENSCAN (Burge and Karlin 1997), the annotated proteins and the nucleotides were then applied to search against GenBank and the Gypsy Database which is an open and well collected database of LTR retrotransposons, retroviruses and pararetroviruses (Llorens et al. 2011).

### Estimation of Copy Numbers of Pararetrovirus in the Barley Genome

The representative pararetrovirus was used as the query to screen the barley genome sequences using the Repeat masker software (http://www.repeatmasker.org). The program was run using the default parameters but ‘nolow’ option. In addition, a cut-off score of ≥ 250 was set. As plant pararetroviruses share certain sequence similarity with LTR retrotransposons at the polymerase-encoding region, to avoid overestimation of pararetrovirus and eliminate the LTR retrotransposon sequences, all output reads were then manually inspected and only the sequences with high confidence (the score > 1000 and the sizes > 150 bp) were accounted.

### Identification of Nested Transposons and Estimation of Insertion Times of LTR Retrotransposons

The RepeatMasker output file of Hvu-EPRVs was manually curated and the sequences that interrupted the Hvu-EPRV elements were extracted and used to search against the barley repeat database (Dongying Gao, unpublished) to identify the nested transposons, the complete TEs were defined by the terminal repeats and the target site duplications (TSDs). To estimate the insertion time of LTR retrotransposons, all intact nested LTR retrotransposons were extracted and conducted BLASTN2 comparisons for determining the exact LTR boundaries of each retroelement. The two LTRs of each elements were aligned and used to estimate the K value (average number of substitutions per aligned site) with the Kimura-2 parameter using MEGA11 (Tamura et al. 2021). The insertion times (T) of LTR retrotransposons were calculated based on the formula: T=K/2r where r represents the average substitution rate of 1.3×10^−8^ substitutions per synonymous site per year.

### Definition of Orthologous EPRVs

One-Kb flanking region for each side of a complete Hvu-EPRV sequence or a tandemly organized Hvu-EPRV block containing complete Hvu-EPRV(s) in the barley reference genome (Morex-V3) was extracted, merged and used to conduct BLASTN search against the published genomes of cultivated and wild barley deposited in GenBank. The significant hits were manually inspected, and the orthologous sequences were defined. For the queries which showed multiple significant hits in the relative genomes, the best hits with highest sequence identity and lowest E-value were considered as the potential orthologous sequences, and their genomic positions were inspected to further confirm the orthologous relationships. In addition, the complete Hvu-EPRVs and their 2-Kb flanking (1-Kb for each side) were extracted and used to compare their orthologous sequences in other genomes with the online mVISTA tool (Frazer et al. 2004) to further confirm the presence or orthologous Hvu-EPRVs.

### Construction of Phylogenetic Tree

The conserved RT protein domains of plant pararetroviruses and the four outgroup LTR retrotransposons were aligned using ClustalW program with default options. The alignments of multiple sequences were applied to build phylogenetic trees with MEGA11 software (Tamura et al. 2021) using the neighbor-joining method. The phylogenetic analysis was evaluated by 500 bootstrap replicate sampling, and the phylogenetic tree of conserved RT proteins was uploaded to the Interactive Tree Of Life website (iTOL: https://itol.embl.de) (Letunic and Bork 2021) and visualized.

## Supplementary Material

Supplementary material is available at the journal’s online.

## Acknowledgments

I would like to thank Drs. Ning Jiang and Noelle Anglin for the valuable comments. The author declares no conflict of interests. The author also declares that the research was conducted in the absence of any commercial or financial relationships that could be construed as a potential conflict of interest. Mention of trade names or commercial products in this publication is solely for the purpose of providing specific information and does not imply recommendation or endorsement by the United States Department of Agriculture.

## Funding

This research was supported by the U.S. Department of Agriculture, Agricultural Research Service.

## Notes

### Competing Interest Statement

The authors have declared no competing interest.

