## Supplementary files for "A Novel Genus of Endogenous Pararetroviruses with Long Terminal Repeats in Grasses"

Supplementary Table S1. A list of homologous sequences of Hvu-EPRV in plant genomes

| EPRVs | Plants | Locations | Long terminal repeats |
|---|---|---|---|
| Nth-EPRV | Nymphaea thermarum | JAANDH010000884.1:21955-30373 | |
| Aca-EPRV | Aristolochia californica | JARWKL010000008.1:37924160-37934326 | |
| Ach-EPRV | Annona cherimola | JASFXW010000158.1:48233-52902 | |
| Ako-EPRV | Amorphophallus konjac | JAHNEJ010000005.1:569187302-569180718 | |
| Abr-EPRV | Acanthochlamys bracteata | JAFKAK010000819.1:5964854-5972887 | |
| Dal-EPRV | Dioscorea alata | CZHE02025868.1:1-6356 | |
| Ata-EPRV | Aegilops tauschii | MCGU01039594.1:89771-99574 | X |
| Amy-EPRV | Alopecurus myosuroides | CASDCE010000002.1:66236157-66227186 | X |
| Dgl-EPRV | Dactylis glomerata | QXEO01001393.1:503-8148 | |
| Lmu-EPRV | Lolium multiflorum | QGDF01006281.1:661657-650277 | X |
| Pan-EPRV | Poa annua | JAPMLF010000008.1:33646581-33637307 | X |
| Pte-EPRV | Puccinellia tenuiflora | QRDG01001467.1:243135-233646 | X |
| Sce-EPRV | Secale cereale | JADQCU010000004.1:144408526-144418097 | X |
| Sca-EPRV | Stipa capillata | JAGXJF010003582.1:840409-844846 | |
| Tel-EPRV | Thinopyrum elongatum | JAAAXO010000002.1:185729704-185738570 | X |
| Tur-EPRV | Triticum urartu | MKGO01000007.1:82207146-82198067 | X |
| Tae-EPRV | Triticum aestivum | NC_057808.1:410525919-410516272 | X |
| Hsp-EPRV | Hordeum spontaneum | OX460006 :675583240-675573790 | X |
| Hma-EPRV | Hordeum marinum | JAAAWK010000004: 433987176-433978012 | X |
| Asa-EPRV | Avena sativa | JAJTTR010000001.1:494623391-494632755 | X |
| Aat-EPRV | Avena atlantica | CAJUWN010002083.1:37011-27714 | X |
| Bdi-EPRV | Brachypodium distachyon | AC276546.1:22533-31795 | X |
| Ase-EPRV | Alloteropsis semialata | QPGU01000681.1:56089892-56098224 | |
| Bde-EPRV | Bothriochloa decipiens | JALGXP010006490.1:356-2758 | |
| Caq-EPRV | Coix aquatica | RZMS01000005.1:5273439-5278631 | |
| Dex-EPRV | Digitaria exilis | JACEFO010002248.1:183743-189609 | |
| Mfl-EPRV | Miscanthus floridulus | JAAIKP010000018.1:47455441-47462511 | |
| Pvi-EPRV | Panicum virgatum | JABWAI010000415.1:1576176-1586169 | |
| Pno-EPRV | Paspalum notatum | JAJQMO010000001.1:17463495-17468777 | |
| Ttr-EPRV | Themeda triandra | JAOPFP010038805.1:299-6106 | |
| Sit-EPRV | Setaria italica | NC_028450.1:27758668-27753616 | |
| Dla-EPRV | Dendrocalamus latiflorus | JACBGG010000025.1:6805958-6815115 | X |
| Ped-EPRV | Phyllostachys edulis | WJQQ01000006.1:3717443-3721738 | |
| Rdi-EPRV | Raddia distichophylla | SPJY01001759.1:2400-3911 | |
| Ecu-EPRV | Eragrostis curvula | RWGY01000643.1:210336-201697 | X |
| Ein-EPRV | Eleusine indica | JARKIL010000001.1:4783788-4790770 | X |
| San-EPRV | Streptochaeta angustifolia | JAHBRV010000035.1:4330519-4338496 | |
| Pau-EPRV | Phragmites australis | JAGFME010011185.1:8044-15956 | |
| Ome-EPRV | Oryza meridionalis | LONC01002787.1:41858-49492 | |
| Osa-EPRV | Oryza sativa | JACXSC010000006.1:3406043-3413697 | |
| Cpa-EPRV | Carex parvula | JAJGRW010005178.1:190946-198316 | |
| Ces-EPRV | Cyperus esculentus | JASGLE010000035.1:3314591-3321882 | |
| Lsy-EPRV | Luzula sylvatica | CAMPEH010000027.1:304956-312002 | |
| Pra-EPRV | Puya raimondii | JABWPP010000049.1:5274215-5279362 | |
| Aco-EPRV | Ananas comosus | BPUW01000338.1:460624-462855 | |
| Mit-EPRV | Musa itinerans | LVTN01000999.1:99481-102864 | |
| Wvi-EPRV | Wurfbainia villosa | JAKLTH010000299.1:3293694-3301481 | |
| Cla-EPRV | Costus lasius | JANVAS010000059.1:3672089-3680638 | |
| Min-EPRV | Macadamia integrifolia | UZVR01000056.1:217958-229702 | |
| Bco-EPRV | Beta corolliflora | CANSZD010000433.1:1888575-1905026 | |

| | | |
|---|---|---|
| Vmy-EPRV | Vaccinium myrtillus | JADGMU010000008.1:22927414-22931166 |
| Car-EPRV | Coffea arabica | RHJU01000019.1:25215698-25218447 |
| Oeu-EPRV | Olea europaea | JAKWBP010000046.1:34222283-34225722 |
| Fdi-EPRV | Fraxinus dipetala | CADEPD010004301.1:264-3824 |
| Lba-EPRV | Lycium barbarum | JAHTKO010000014.1:794463-802059 |
| Sar-EPRV | Solanum arcanum | JAPXVU010000009.1:60417927-60426870 |
| Sve-EPRV | Solanum verrucosum | FXZR01005138.1:4874-12617 |
| Pgr-EPRV | Platycodon grandiflorus | SPEA01004552.1:39267-44384 |
| Afo-EPRV | Anethum foeniculum | PHNY01262050.1:13082-17818 |
| Iaq-EPRV | Ilex aquifolium | CATOGZ010000263.1:316637-323365 |
| Ltu-EPRV | Lathyrus tuberosus | ALAZF010000100.1:1865543-1872900 |
| Vun-EPRV | Vigna unguiculata | BLZF01002321.1:888486-896042 |
| Msy-EPRV | Malus sylvestris | CAJZLB010000703.1:92165-100714 |
| Fmi-EPRV | Ficus microcarpa | JAINRI010000006.1:1400712-1403243 |
| Hlu-EPRV | Humulus lupulus | JALDWI010005821.1:91055-98617 |
| Rru-EPRV | Rhamnella rubrinervis | VOIH02000003.1:5288937-5296697 |
| Twi-EPRV | Tripterygium wilfordii | JAAARO010000012.1:12932904-12941370 |
| Qsu-EPRV | Quercus suber | PKMF03021175.1:636602-637912 |
| Blo-EPRV | Begonia loranthoides | JAJTVF010006050.1:4393-5967 |
| Epu-EPRV | Euphorbia pulcherrima | CANTUS010000012.1:2972111-2982415 |
| Cmi-EPRV | Combretum micranthum | JAMXBD010000206.1:1157861-1167510 |
| Sle-EPRV | Shorea leprosula | BPVZ01000017.1:883814-888884 |
| Ain-EPRV | Azadirachta indica | JAGQDM010000003.1:16152832-16156257 |
| Car-EPRV | Citropsis articulata | JASUUH010000200.1:3133147-3139117 |
| Pac-EPRV | Polystichum acrostichoides | JAOYMV010455690.1:1-3136 |
| Dma-EPRV | Dryopteris marginalis | JAODKW010142212.1:1-3985 |
| Cla-EPRV | Cunninghamia lanceolata | BSBN01000768.1:4961080-4967662 |
| Tch-EPRV | Taxus chinensis | JAHRHJ020000002.1:9901846-9905920 |
| Pla-EPRV | Pinus lambertiana | LMTP010341120.1:1301-7379 |
| Gbi-EPRV | Ginkgo biloba | JANKJI010143559.1:1-1256 |
| Gmo-EPRV | Gnetum montanum | MNCI01023097.1:812626-819695 |

Supplementary Table S2. 37 reported retroelements that are used for phylogenetic analysis

| Pararetrovirus | Classification | GenBank or (Literatures) |
|---|---|---|
| Ty3 | LTR retrotransposon | M34549 |
| Gypsy | LTR retrotransposon | M12927 |
| Jinling | LTR retrotransposon | DQ445619 |
| FRetro3 | LTR retrotransposon | GU369679 |
| Citrus yellow mosaic virus (CiYMV) | Badnavirus | NC_003382 |
| Banana streak GF virus (BSGFV) | Badnavirus | NC_007002 |
| Cacao swollen shoot virus (CSSV) | Badnavirus | NP_041734 |
| Kalanchoe top-spotting virus (KTSV) | Badnavirus | NC_004540 |
| Cauliflower mosaic virus (CaMV) | Caulimovirus | NC_001497 |
| Strawberry vein banding virus (SVBV) | Caulimovirus | NC_001725 |
| Carnation etched ring virus (CERV) | Caulimovirus | NC_003498 |
| Horseradish latent virus (HRLV) | Caulimovirus | JX429923.1 |
| Angelica bushy stunt virus (AnBSV) | Caulimovirus | AMN10080.1 |
| Cassava vein mosaic virus (CSVMV) | Cavemovirus | NC_001648 |
| Dioscorea nummularia-associated virus (DNAV) | Dioscovirus | YP_009553219 |
| Petunia vein clearing virus (PVCV) | Petuvirus | NC_001839 |
| Rose yellow vein virus (RYVV) | Rosadnavirus | YP_007761644 |
| Tobacco vein clearing virus (TVCV) | Solendovirus | AF190123 |
| Peanut chlorotic streak virus (PCSV) | Soymovirus | NC_001634 |
| Soybean chlorotic mottle virus (SbCMV) | Soymovirus | NC_001739 |
| Blueberry red ringspot virus (BRRV) | Soymovirus | NC_003138 |
| Rice tungro bacilliform virus (RTBV) | Tungrovirus | AJ314596 |
| Dioscorea nummularia-associated virus | | CANXTL010000001 |
| Nicotiana tabacum pararetrovirus (NtEPRV) | | (Jakowitsch et al. 1999) |
| Citrus endogenous pararetrovirus (CiERV) | | (Roy et al. 2014) |
| Piper nigrum endogenous virus (PNEV1) | | (Bhat et al. 2022) |
| Ah_ParaRetro1 | | (Bertioli et al. 2019) |
| Ah_ParaRetro2 | | (Bertioli et al. 2019) |
| Fernendo2_Dipteris | | (Vassilieff et al. 2022) |
| Florendovirus_Nsyl | Florendovirus | (Vassilieff et al. 2022) |
| Gymnendovirus_1 | Gymnendovirus | Vassilieff et al. 2022) |
| Petuviruslike_Csine | Xendovirus | (Vassilieff et al. 2022) |
| Xendovirus_Grai | Xendovirus | (Vassilieff et al. 2022) |
| Yendovirus_Anaco | Yendovirus | (Vassilieff et al. 2022) |
| Fvesca | | (Vassilieff et al. 2022) |
| CitIch-033 | Wendovirus | (de Tomás and Vicient 2022) |
| Slyc_Paraseq_1592 | | (Valli et al. 2023) |

Supplementary Table S3. The distributions of Hvu-EPRVs at eight genomic loci in 85 cultivated and wild genomes

| Genomes | Species | **I** | **II** | **III** | **IV** | **V** | **VI** | **VII** | **VIII** | References |
|---|---|---|---|---|---|---|---|---|---|---|
| MorexV3 | *H. vulgare* | p | p | p | p | p | p | p | p | Jayakodi et al. 2024 |
| HOR_12184_BPGv2 | *H. vulgare* | p | p | p | a | a | a | p | p | Jayakodi et al. 2024 |
| HOR_8117_BPGv2 | *H. vulgare* | p | p | a | p | a | a | p | p | Jayakodi et al. 2024 |
| HOR_2830_BPGv2 | *H. vulgare* | p | p | a | p | a | a | a | p | Jayakodi et al. 2024 |
| HOR_13821_BPGv2 | *H. vulgare* | p | a | a | p | a | a | p | p | Jayakodi et al. 2024 |
| HOR_2180_BPGv2 | *H. vulgare* | p | a | p | a | p | a | a | p | Jayakodi et al. 2024 |
| HOR_8148_BPGv2 | *H. vulgare* | p | a | a | p | a | a | p | p | Jayakodi et al. 2024 |
| Hulless_Barley_ass.V2 | *H. vulgare* | p | a | a | a | p | p | p | a | Zeng et al. 2020 |
| HOR_7552_BPGv2 | *H. vulgare* | p | a | a | a | p | p | p | a | Jayakodi et al. 2024 |
| HOR_495_BPGv2 | *H. vulgare* | a | p | p | a | p | a | a | p | Jayakodi et al. 2024 |
| Golden_Melon_BPGv2 | *H. vulgare* | a | p | a | p | a | p | a | p | Jayakodi et al. 2024 |
| HOR_9043_BPGv2 | *H. vulgare* | p | a | a | p | a | a | a | p | Jayakodi et al. 2024 |
| HOR_6220_BPGv2 | *H. vulgare* | p | a | a | p | a | a | a | p | Jayakodi et al. 2024 |
| HOR_7172_BPGv2 | *H. vulgare* | p | a | a | a | p | p | a | a | Jayakodi et al. 2024 |
| HOR_13663_BPGv2 | *H. vulgare* | a | p | a | a | a | a | p | p | Jayakodi et al. 2024 |
| HOR_3081_BPGv2 | *H. vulgare* | a | p | a | a | a | a | p | p | Jayakodi et al. 2024 |
| HOR_21256_BPGv2 | *H. vulgare* | a | a | p | p | a | a | a | p | Jayakodi et al. 2024 |
| HID380_BPGv2 | *H. vulgare* | a | a | a | p | a | p | a | p | Jayakodi et al. 2024 |
| HOR_14061_BPGv2 | *H. vulgare* | a | a | a | p | a | a | p | p | Jayakodi et al. 2024 |
| HOR_21599_BPGv2 | *H. vulgare* | a | a | a | p | a | a | p | p | Jayakodi et al. 2024 |
| Du_Li_Huang_ZDM01467_BPGv2 | *H. vulgare* | a | a | a | a | p | p | p | a | Jayakodi et al. 2024 |
| 10TJ18_BPGv2 | *H. vulgare* | p | a | a | a | p | a | a | a | Jayakodi et al. 2024 |
| HOR_10350_BPGv2 | *H. vulgare* | p | a | a | p | a | a | a | a | Jayakodi et al. 2024 |
| HOR_10892_BPGv2 | *H. vulgare* | a | p | a | a | a | a | p | a | Jayakodi et al. 2024 |
| Aizu6_BPGv2 | *H. vulgare* | a | p | a | a | a | a | a | p | Jayakodi et al. 2024 |
| HOR_4224_BPGv2 | *H. vulgare* | a | p | a | a | a | a | a | p | Jayakodi et al. 2024 |
| Stirling | *H. vulgare* | a | a | p | a | a | a | a | p | Hu et al. 2023 |
| Maximus_BPGv2 | *H. vulgare* | a | a | p | a | a | a | a | p | Jayakodi et al. 2024 |
| HOR_21322_BPGv2 | *H. vulgare* | a | a | a | p | a | a | p | a | Jayakodi et al. 2024 |
| HOR_1702_BPGv2 | *H. vulgare* | a | a | a | p | a | a | a | p | Jayakodi et al. 2024 |
| Golden_Promise_BPGv2 | *H. vulgare* | a | a | a | p | a | a | a | p | Jayakodi et al. 2024 |
| RGT_Planet_BPGv2 | *H. vulgare* | a | a | a | p | a | a | a | p | Jayakodi et al. 2024 |
| Barke_BPGv2 | *H. vulgare* | a | a | a | p | a | a | a | p | Jayakodi et al. 2024 |
| HOR_13594_BPGv2 | *H. vulgare* | a | a | a | p | a | a | a | p | Jayakodi et al. 2024 |
| Bonus_BPGv2 | *H. vulgare* | a | a | a | p | a | a | a | p | Jayakodi et al. 2024 |
| HOR_14273_BPGv2 | *H. vulgare* | a | a | a | p | a | a | a | p | Jayakodi et al. 2024 |
| HOR_9972_BPGv2 | *H. vulgare* | a | a | a | a | p | a | p | a | Jayakodi et al. 2024 |
| HOR_2779_BPGv2 | *H. vulgare* | a | a | a | a | a | p | p | a | Jayakodi et al. 2024 |
| HOR_7385_BPGv2 | *H. vulgare* | a | a | a | a | a | p | p | a | Jayakodi et al. 2024 |
| Igri_BPGv2 | *H. vulgare* | a | a | a | a | a | a | p | p | Jayakodi et al. 2024 |
| OUN333_BPGv2 | *H. vulgare* | p | a | a | a | a | a | a | a | Jayakodi et al. 2024 |
| HOR_13942_BPGv2 | *H. vulgare* | p | a | a | a | a | a | a | a | Jayakodi et al. 2024 |
| Clipper | *H. vulgare* | a | a | a | p | a | a | a | a | Hu et al. 2023 |

| | | | | | | | | | | |
|---|---|---|---|---|---|---|---|---|---|---|
| Foma_BPGv2 | *H. vulgare* | a | a | a | p | a | a | a | a | Jayakodi et al. 2024 |
| HOR_19184_BPGv2 | *H. vulgare* | a | a | a | p | a | a | a | a | Jayakodi et al. 2024 |
| HOR_18321_BPGv2 | *H. vulgare* | a | a | a | a | a | a | p | a | Jayakodi et al. 2024 |
| Hockett_BPGv2 | *H. vulgare* | a | a | a | a | a | a | a | p | Jayakodi et al. 2024 |
| Chikurin_Ibaraki1_BPGv2 | *H. vulgare* | a | a | a | a | a | a | a | p | Jayakodi et al. 2024 |
| Akashinriki_BPGv2 | *H. vulgare* | a | a | a | a | a | a | a | p | Jayakodi et al. 2024 |
| AAC_Synergy_1.0 | *H. vulgare* | a | a | a | ? | a | a | a | p | Xu et al. 2021 |
| HOR_21595_BPGv2 | *H. vulgare* | a | a | a | a | a | a | a | p | Jayakodi et al. 2024 |
| HOR_14121_BPGv2 | *H. vulgare* | ? | a | a | a | a | a | a | p | Jayakodi et al. 2024 |
| CI5791v1 | *H. vulgare* | a | a | a | a | a | a | a | p | Clare et al. 2024 |
| HOR_3365_BPGv2 | *H. vulgare* | a | a | a | a | a | a | a | p | Jayakodi et al. 2024 |
| HOR_3474_BPGv2 | *H. vulgare* | a | a | a | a | a | a | a | p | Jayakodi et al. 2024 |
| Chiba_ZDM02064_BPGv2 | *H. vulgare* | a | a | a | a | a | a | a | p | Jayakodi et al. 2024 |
| Bowman_BPGv2 | *H. vulgare* | a | a | a | a | a | a | a | P | Jayakodi et al. 2024 |
| HTX_v1 | *H. vulgare* | a | a | a | a | a | a | a | a | Jiang et al. 2022 |
| HOR_1168_BPGv2 | *H. vulgare* | a | a | a | a | a | a | a | a | Jayakodi et al. 2024 |
| HOR_10096_BPGv2 | *H. vulgare* | a | a | a | a | a | a | a | a | Jayakodi et al. 2024 |
| HOR_12541_BPGv2 | *H. vulgare* | a | a | a | a | a | a | a | a | Jayakodi et al. 2024 |
| FT11_B1K_04_12_BPGv2 | *H. spontaneum* | p | a | a | a | a | p | p | a | Jayakodi et al. 2024 |
| FT880_BPGv2 | *H. spontaneum* | ? | a | a | a | a | a | p | p | Jayakodi et al. 2024 |
| WBDC_207_BPGv2 | *H. spontaneum* | p | a | a | a | a | a | p | a | Jayakodi et al. 2024 |
| WBDC_184_BPGv2 | *H. spontaneum* | p | a | a | a | a | a | p | a | Jayakodi et al. 2024 |
| WBDC_348_BPGv2 | *H. spontaneum* | p | a | a | a | a | p | a | a | Jayakodi et al. 2024 |
| Zhang lab wild barley-ZWY2020 | *H. spontaneum* | ? | a | a | a | p | a | p | a | Liu et al. 2020 |
| HID357_BPGv2 | *H. spontaneum* | p | a | a | a | a | a | a | a | Jayakodi et al. 2024 |
| ASM2978338v1-EC-N1 | *H. spontaneum* | a | a | a | a | a | p | a | a | Pan et al. 2023 |
| FT286_BPGv2 | *H. spontaneum* | a | a | a | a | a | a | p | a | Jayakodi et al. 2024 |
| FT67_BPGv2 | *H. spontaneum* | a | a | a | a | a | a | p | a | Jayakodi et al. 2024 |
| WBDC_078_BPGv2 | *H. spontaneum* | a | a | a | a | a | a | p | a | Jayakodi et al. 2024 |
| WBDC_103_BPGv2 | *H. spontaneum* | a | a | a | a | a | a | p | a | Jayakodi et al. 2024 |
| ASM2978261v1-EC-S1 | *H. spontaneum* | a | a | a | a | a | a | p | a | Pan et al. 2023 |
| FT333_BPGv2 | *H. spontaneum* | a | a | a | a | a | a | a | p | Jayakodi et al. 2024 |
| FT262_BPGv2 | *H. spontaneum* | a | a | a | a | a | a | a | a | Jayakodi et al. 2024 |
| HID055_BPGv2 | *H. spontaneum* | a | a | a | a | a | a | a | a | Jayakodi et al. 2024 |
| WBDC_199_BPGv2 | *H. spontaneum* | a | a | a | a | a | a | a | a | Jayakodi et al. 2024 |
| FT144_BPGv2 | *H. spontaneum* | a | a | a | a | a | ? | a | a | Jayakodi et al. 2024 |
| WBDC_349_BPGv2 | *H. spontaneum* | a | a | a | a | a | a | a | a | Jayakodi et al. 2024 |
| WBDC_237_BPGv2 | *H. spontaneum* | a | a | a | a | a | a | a | a | Jayakodi et al. 2024 |
| FT628_BPGv2 | *H. spontaneum* | a | a | a | a | a | a | a | a | Jayakodi et al. 2024 |
| WBDC_133_BPGv2 | *H. spontaneum* | a | a | a | a | a | a | a | a | Jayakodi et al. 2024 |
| HID101_BPGv2 | *H. spontaneum* | a | a | a | a | a | a | a | a | Jayakodi et al. 2024 |
| HID249_BPGv2 | *H. spontaneum* | a | a | a | a | a | a | a | a | Jayakodi et al. 2024 |

Note:1) "p" means presence, "a" means absence, and "?" means unknown as some flanking regions were not detected or highly repetitive; The locations of Hvu-EPRVs in Morex: I (LR890097: 8896806-8906251), II (LR890098: 345974868-345984153), III (LR890099: 406022753-406057905), IV (LR890099: 465067318-465097321), V (LR890099: 491070135-491087673), VI (LR890101: 35626967-35636343), VII (LR890101: 88225996-88235310) and VIII (LR890101: 96416978-96454773).

Supplementary Table S4. A list of Hvu-EPRV related ESTs/cDNAs from 17 grasses

| GenBank accession | E value | Species |
|---|---|---|
| GT766930.1 | 2E-66 | Brachypodium distachyon |
| GT853149.1 | 3E-39 | Brachypodium distachyon |
| GT853150.1 | 6E-36 | Brachypodium distachyon |
| GT855714.1 | 9.00E-34 | Brachypodium distachyon |
| GT855713.1 | 9.00E-34 | Brachypodium distachyon |
| GT845740.1 | 5.00E-31 | Brachypodium distachyon |
| GT845739.1 | 2.00E-28 | Brachypodium distachyon |
| GT766929.1 | 8.00E-22 | Brachypodium distachyon |
| ES302538.1 | 0 | Cynodon dactylon |
| HO129612.1 | 2E-99 | Dactylis glomerata |
| HO161807.1 | 4E-63 | Dactylis glomerata |
| HO130804.1 | 7E-48 | Dactylis glomerata |
| HO147785.1 | 5E-43 | Dactylis glomerata |
| HO167501.1 | 5E-43 | Dactylis glomerata |
| HO148974.1 | 2E-41 | Dactylis glomerata |
| HO149737.1 | 3E-39 | Dactylis glomerata |
| HO122542.1 | 7.00E-23 | Dactylis glomerata |
| HO123120.1 | 1.00E-20 | Dactylis glomerata |
| FF596965.1 | 4.00E-25 | Elymus wawawaiensis/Elymus lanceolatus |
| GO887646.1 | 6E-112 | Festuca pratensis |
| GO887611.1 | 6E-112 | Festuca pratensis |
| BQ755041.1 | 2.00E-23 | Hordeum vulgare |
| GT040988.1 | 0 | Lolium arundinaceum |
| DT706952.1 | 7.00E-29 | Lolium arundinaceum |
| HS376650.1 | 1E-56 | Oryza longistaminata |
| HS370321.1 | 6E-55 | Oryza longistaminata |
| HS340308.1 | 2E-47 | Oryza longistaminata |
| HS371896.1 | 2.00E-30 | Oryza longistaminata |
| HS386657.1 | 8.00E-22 | Oryza longistaminata |
| CB884243.1 | 6.00E-17 | Oryza minuta |
| CK038670.1 | 2E-157 | Oryza sativa |
| CT862892.1 | 6E-125 | Oryza sativa |
| CA763970.2 | 8E-111 | Oryza sativa |
| CK056718.1 | 2E-100 | Oryza sativa |
| CT849564.1 | 3E-59 | Oryza sativa |
| FG946986.1 | 6E-55 | Oryza sativa |

| | | |
|---|---|---|
| CI036069.1 | 6E-49 | Oryza sativa |
| CK045057.1 | 2E-41 | Oryza sativa |
| AK062971 | 3.00E-40 | Oryza sativa |
| CF310066.1 | 9.00E-28 | Oryza sativa |
| CF310369.1 | 2.00E-24 | Oryza sativa |
| CF960273.1 | 5.00E-24 | Oryza sativa |
| CX105740.1 | 3.00E-20 | Oryza sativa |
| JG866652.1 | 2E-47 | Panicum virgatum |
| JG965617.1 | 1.00E-20 | Panicum virgatum |
| JG878483.1 | 1.00E-20 | Panicum virgatum |
| JG865619.1 | 1.00E-19 | Panicum virgatum |
| JG901705.1 | 4.00E-19 | Panicum virgatum |
| FL710451.1 | 6.00E-17 | Panicum virgatum |
| GD012924.1 | 1.00E-13 | Panicum virgatum |
| FF361881.1 | 1E-76 | Pseudoroegneria spicata |
| FF361882.1 | 1.00E-26 | Pseudoroegneria spicata |
| FF349483.1 | 6.00E-17 | Pseudoroegneria spicata |
| CA094702.1 | 1.00E-25 | Saccharum hybrid cultivar |
| EC325393.1 | 2E-49 | Saccharum hybrid |
| CA117698.1 | 5E-164 | Saccharum hybrid cultivar |
| CA168610.1 | 2.00E-28 | Saccharum hybrid cultivar |
| CA171713.1 | 1.00E-26 | Saccharum hybrid cultivar |
| CA266811.1 | 1.00E-25 | Saccharum hybrid cultivar |
| CA097829.1 | 1.00E-25 | Saccharum hybrid cultivar |
| CA267242.1 | 2.00E-24 | Saccharum hybrid cultivar |
| CA264653.1 | 2.00E-24 | Saccharum hybrid cultivar |
| CA095975.1 | 2.00E-24 | Saccharum hybrid cultivar |
| CA171062.1 | 5.00E-24 | Saccharum hybrid cultivar |
| CA143105.1 | 8.00E-22 | Saccharum hybrid cultivar |
| CA097826.1 | 8.00E-22 | Saccharum hybrid cultivar |
| CA153576.1 | 2.00E-17 | Saccharum hybrid cultivar |
| CA271879.1 | 8.00E-16 | Saccharum hybrid cultivar |
| BQ478983.1 | 4E-89 | Saccharum officinarum |
| CD204576.1 | 1.00E-13 | Sorghum bicolor |
| CK210164.1 | 0 | Triticum aestivum |
| CV760190.1 | 8E-117 | Triticum aestivum |
| BF484123.1 | 9E-104 | Triticum aestivum |
| CO347755.1 | 4E-83 | Triticum aestivum |
| LU057297.1 | 7E-67 | Triticum aestivum |
| LU004134.1 | 7E-67 | Triticum aestivum |

| | | |
|---|---|---|
| HX063218.1 | 3E-65 | Triticum aestivum |
| AK333052.1 | 4E-54 | Triticum aestivum |
| HX063187.1 | 9E-47 | Triticum aestivum |
| CD917802.1 | 2.00E-24 | Triticum aestivum |
| CJ696218.1 | 3.00E-21 | Triticum aestivum |
| GH724277.1 | 3.00E-20 | Triticum aestivum |
| CJ651881.1 | 1.00E-18 | Triticum aestivum |
| CD910168.1 | 8.00E-16 | Triticum aestivum |
| CA719195.1 | 1.00E-12 | Triticum aestivum |
| CJ537392.1 | 2.00E-11 | Triticum aestivum |
| AJ716778.1 | 4E-114 | Triticum turgidum subsp. durum |

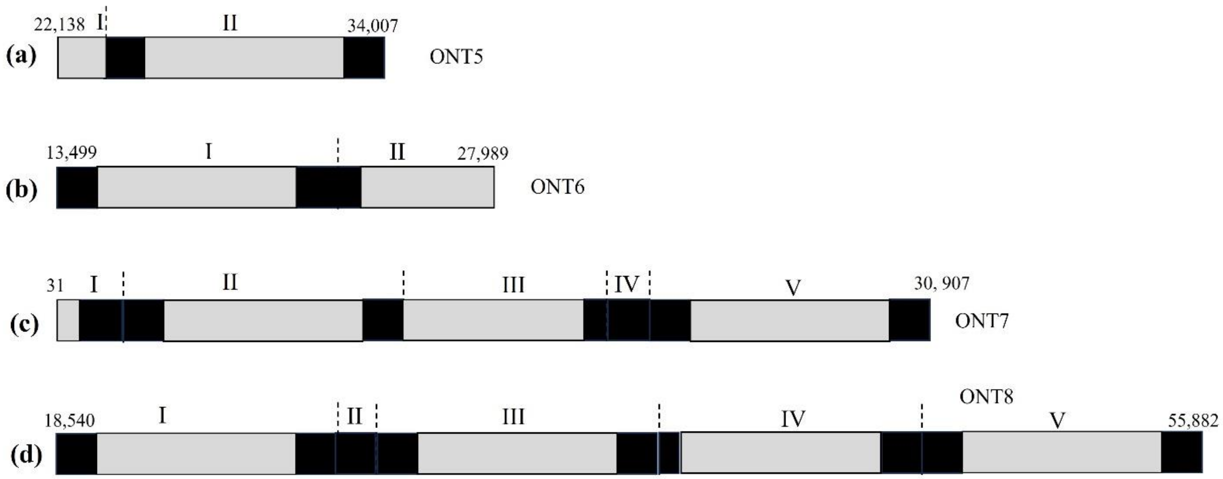

**Supplementary Figure S1. Tandem organizations of Hvu-EPRVs in four barley ONT long-reads.** The black and grey rectangles mean LTDRs and internal regions of Hvu-EPRVs. The different copies of Hvu-EPRVs are indicated by Roman numerals which were defined based on the 9,446-bp reference element.

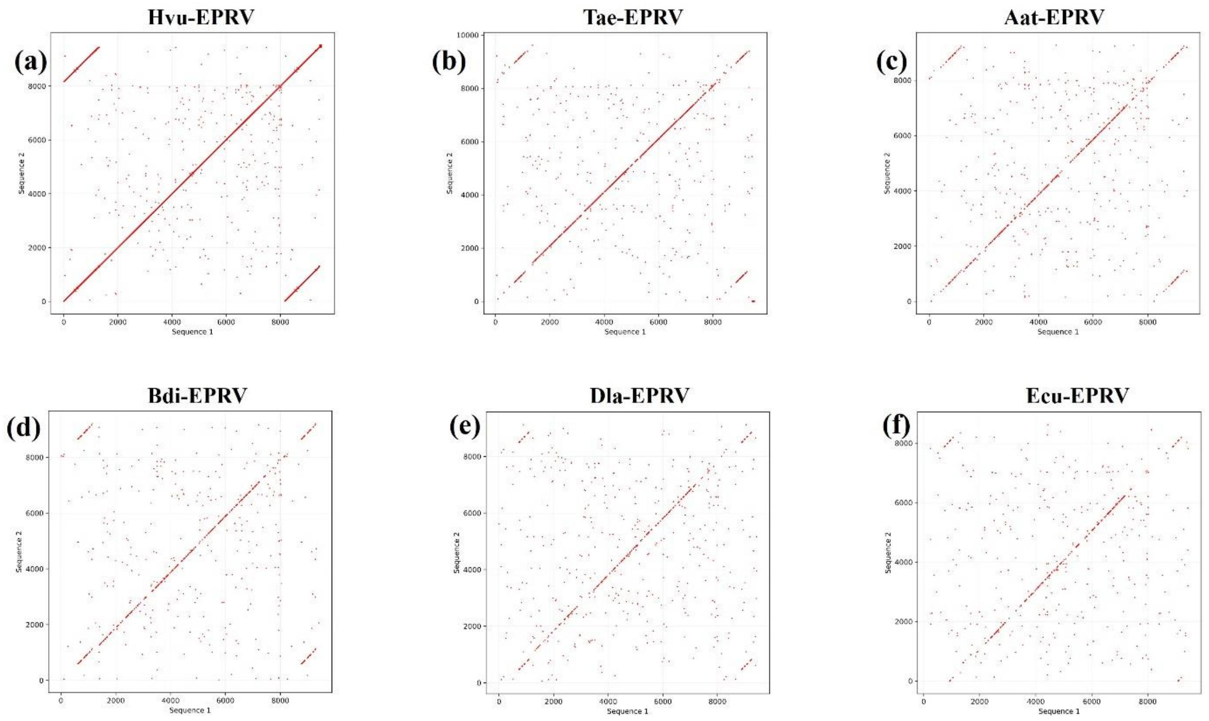

**Supplementary Figure S2. Dot-plot between Hvu-EPRV and Hvu-EPRV (a), Hvu-EPRV and Tae-EPRV in common wheat (b), Hvu-EPRV and Aat-EPRV in diploid oat (c), Hvu-EPRV and Bdi-EPRV in purple false brome (d), Hvu-EPRV and Dla-EPRV in Taiwan giant bamboo (e), Hvu-EPRV and Ecu-EPRV in weeping lovegrass (f). The Dot-plot analysis was conducted with the VectorBuilder online source (https://en.vectorbuilder.com/tool/sequence-dot-plot.html).**

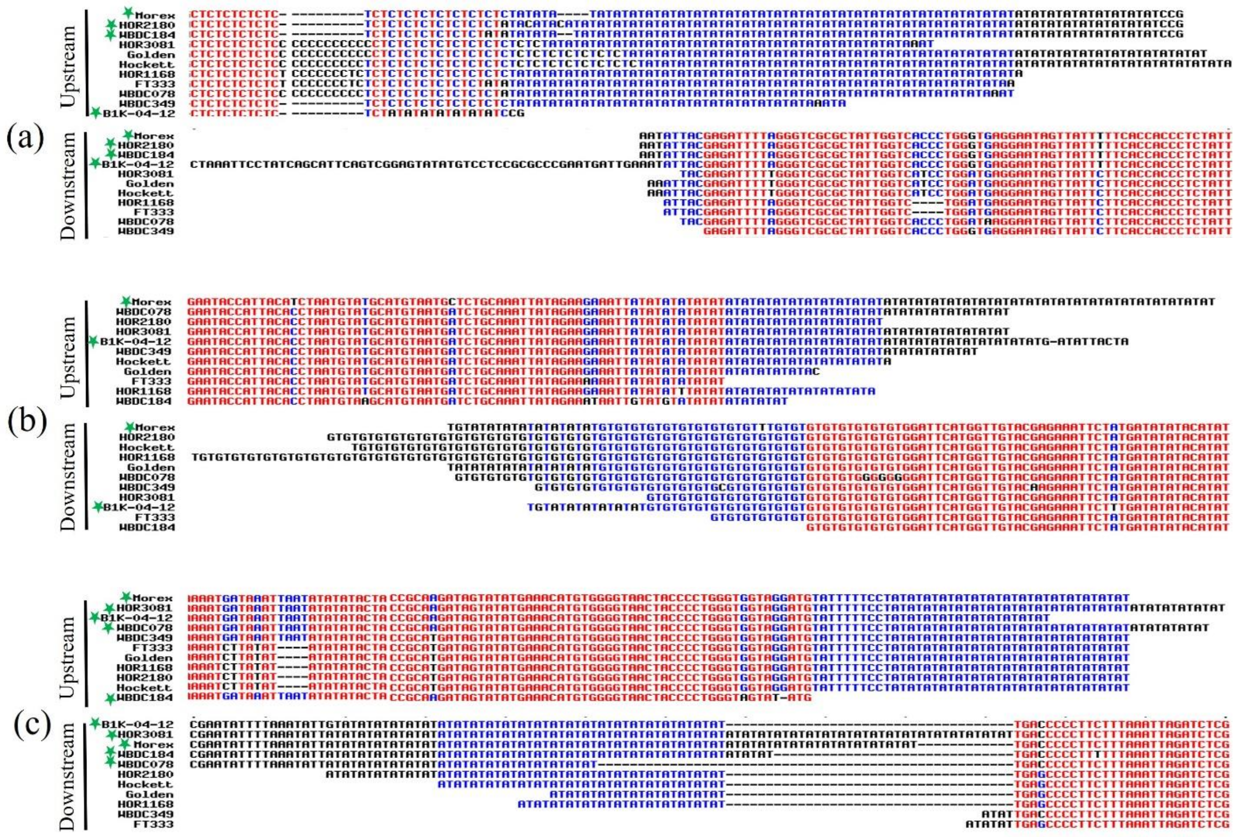

**Supplementary Figure S3. Alignment of 5' (up) and 3' (down) flanking sequences of the Hvu-EPRV in Morex and 10 cultivated and wild barley genomes at the locus I (LR890097: 8896806-8906251, a), VI (LR890101: 35626967-35636343, b) and VII (LR890101: 88225996-88235310, c).** The green stars mean the genotypes containing Hvu-EPRV.

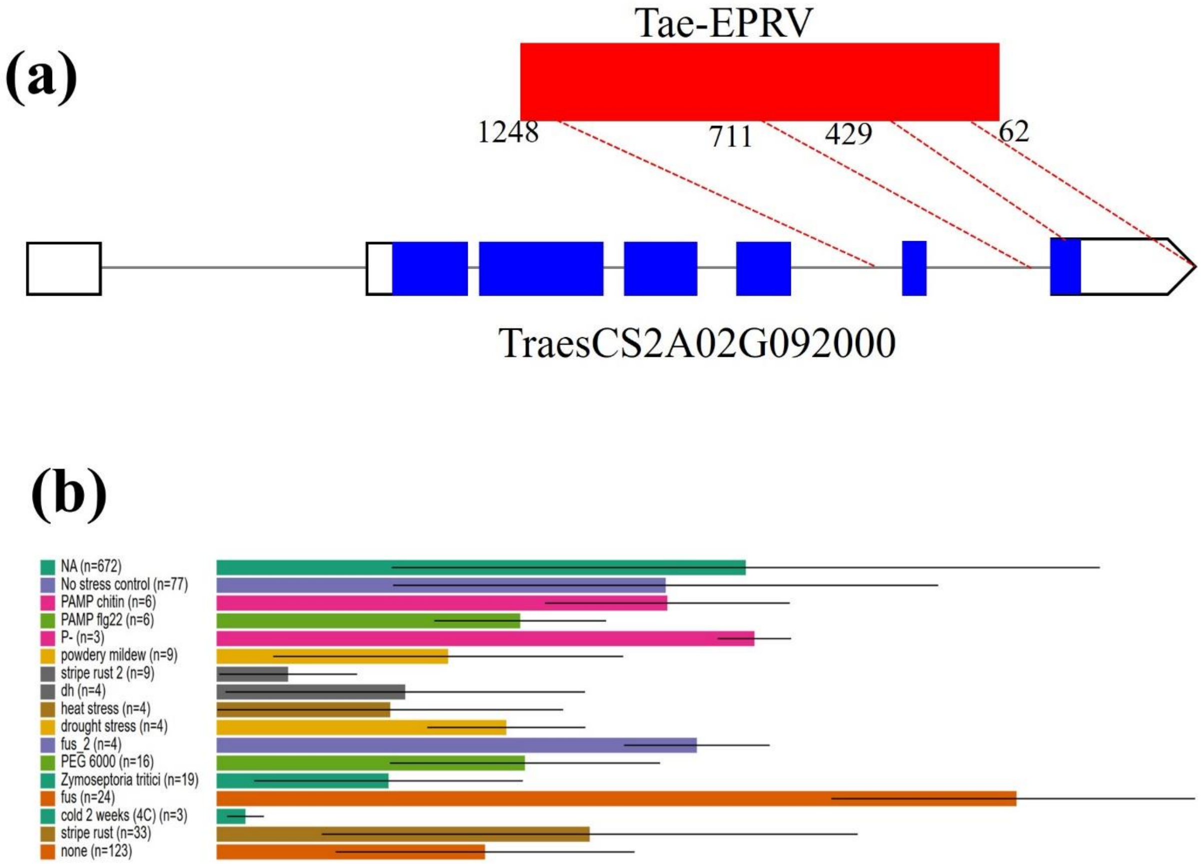

**Supplementary Figure S4. a. The gene structure of *TraesCS2A02G092000* which contains partial LTR sequence of Tae-EPRV**. The blue rectangles mean exons, black lines indicate introns, and the 5' and 3' UTR are represented by white rectangles and white pentagon. The broken lines indicate the two regions of LTR (62-429 bp and 711-1248 bp) that served as genic sequences. **b. The expression of *TraesCS2A02G092000* gene under stresses including inoculations with pathogens.**
